# Unveiling the molecular basis of disease co-occurrence: towards personalized comorbidity profiles

**DOI:** 10.1101/431312

**Authors:** Jon Sánchez-Valle, Hector Tejero, José María Fernández, David Juan, Salvador Capella-Gutiérrez, Fatima Al-Shahrour, Rafael Tabarés-Seisdedos, Vera Pancaldi, Alfonso Valencia

## Abstract

Comorbidity is an impactful medical problem that is attracting increasing attention in healthcare and biomedical research. However, little is known about the molecular processes leading to the development of a specific disease in patients affected by other conditions. We present a disease interaction network inferred from similarities in patients’ molecular profiles, which significantly recapitulates epidemiologically documented comorbidities, providing the basis for their interpretation at a molecular level. Furthermore, expanding on the analysis of subgroups of patients with similar molecular profiles, our approach discovers comorbidity relations not previously described, implicates distinct genes in such relations, and identifies drugs whose side effects are potentially associated to the observed comorbidities.

---

Comorbidity is the tendency for one patient to have an altered risk of developing a second disease when they are already suffering from a specific one. Comorbidity incidence increases with age and has a high impact on life expectancy, which decreases considerably in the presence of a handful of simultaneous diseases ^1^ as is commonly observed in ageing populations ^2^. Additionally, the presence of comorbid conditions has a high economic impact as shown, for example, by the increase of 150% of the cost associated to diabetes for people who are also affected by heart disease ^3^. Thus, it is clear that controlling patient-specific risks of future comorbidities could increase life expectancy and reduce public health expenditure ^4^. In the research area of comorbidity, tens of disease-disease interaction networks have been published since 2007 ^5^, using a variety of data types, such as gene expression profiles ^6^, combination of disease genes and protein-protein interaction networks ^7^, miRNA expression8, the microbiome ^9^, medical claims ^10^, medical records ^11^, human symptoms ^12^, insurance claims ^13^, and mixed information ^14^. The Jensen et al. study considers that patients with the same disease might present different risks of developing secondary diseases based on their medical history ^11^, which can be a consequence of the **existence of different clinical phenotypes within multifaceted conditions as described in chronic obstructive pulmonary disease** ^15^. Therefore, in this study we set out to explore the molecular bases of comorbidity using patients’ transcriptomic profiles to define personalized comorbidity risks. In a previous study based on differential gene expression meta-analyses ^16^, we detected that inversely comorbid Central Nervous System disorders and cancers presented significant overlaps between genes deregulated in opposite directions in the two sets of diseases, providing initial molecular evidence for such comorbidity relations. In this new study we have explored this principle at a different level, calculating differential expression profiles for each patient to reduce samples’ tissue of origin effect, and defining a patient similarity network including over 6,000 patients affected by 133 diseases, including 15 of the top 20 leading causes of death worldwide in 2015 ^17^.

We used patients’ molecular similarity to calculate relative risk interactions between diseases recovering relations that significantly match previously described epidemiological networks. Additionally, we extracted distinct patient-subgroups in most diseases, estimated their relative risk relations and identified subgroups defying general tendencies. Finally, we successfully assigned patients into their corresponding subgroups, and provide proof-of-concept strategies to define personalized comorbidity risk profiles.

## Results

### Diseases’ Molecular Similarity Network

To study patient-specific comorbidities, we collected gene expression data from microarray assays for 6,284 patients suffering from 133 diseases and healthy individuals with 3 phenotypes (smoking, aging, and exercise). We identified differentially expressed genes by comparing each case sample (from now on patient) to all the control samples from the same study. Then, we looked for molecular similarities among patients, based on the significant overlap between the top 500 up-and down-regulated genes ^16^ (Methods, Fig. 1). Studying the molecular similarities between patients based on the expression changes observed in the case sample compared to the controls from the same tissue reduces the tissue of origin effect, as previously described in the inverse comorbidity relations between Alzheimer’s disease (AD) and non-small cell lung cancer (NSCLC) ^18^. Patients with genes deregulated in the same direction (both up-and down-regulated) were connected by a positive interaction, whereas patients showing overlaps between genes deregulated in opposite directions (up-regulated in one patient and down-regulated in the other one and vice versa) were assigned a negative interaction.

**Figure 1.**
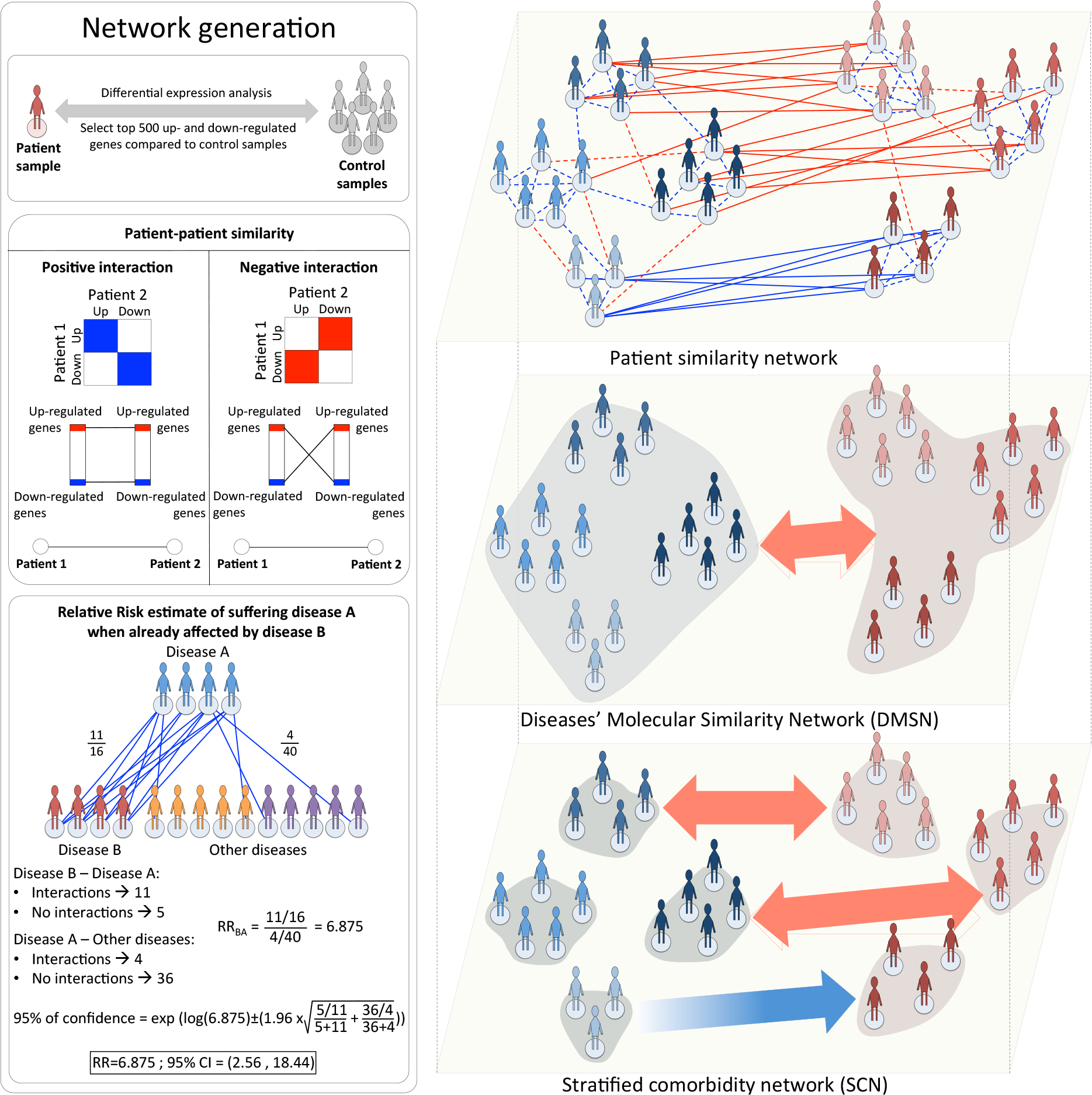
Workflow of the overall study, starting from the differential gene expression analysis, moving to the patient similarity network and the generation of the disease similarity and the stratified comorbidity networks.

We then generated the **Disease’s Molecular Similarity Network (DMSN) calculating positive (pRR) and negative (nRR) relative risks between diseases based on similarities of expression profiles** (Methods) with a confidence interval of 95%. In this sense, a pRR between diseases A and B means that patients with disease A are potentially at a higher risk of developing disease B compared to all the other patients, based on patients’ molecular similarity. On the other hand, a nRR interaction means that patients with disease A are potentially at a significantly lower risk of developing disease B than the rest of the patients in the network. The resulting DMSN is composed of 136 nodes (all diseases and conditions considered in this study) and 5,826 edges (Fig. 2). As expected, most of the RR interactions were pRR (55%), with 24% of them being interactions between diseases from the same ICD9 disease category. Neoplasms were the most connected category.

**Figure 2.**
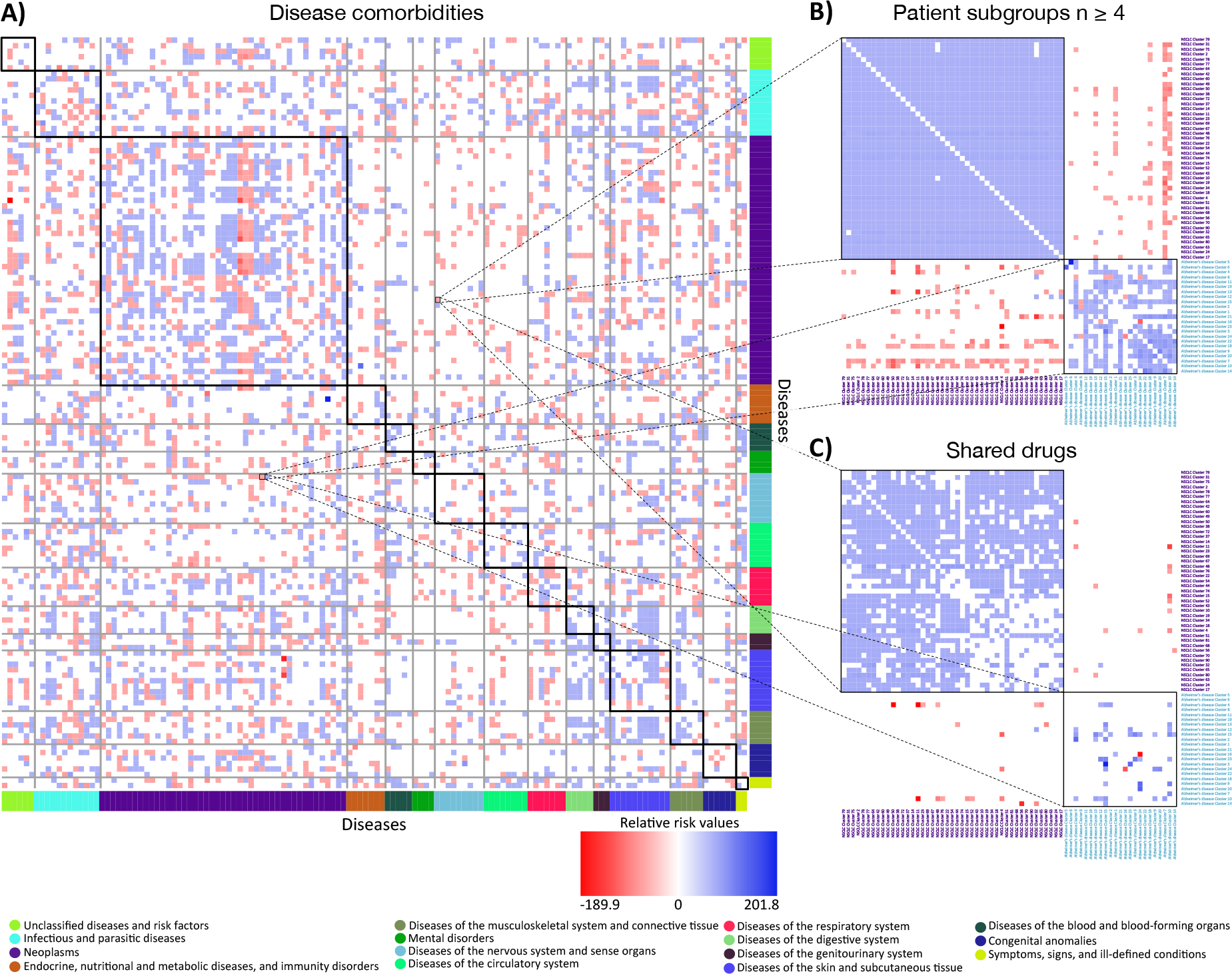
Disease-disease interaction network and Stratified Comorbidity Network. **A)** Heatmap of the disease-disease relative risk interactions. Blue and red squares represent positive and negative relative risks respectively. Intensity of the interactions denote the relative risk values. Relative risk interactions’ directions go from rows to columns. Diseases are colored based on the disease-group they belong to. **B)** Heatmap of the interactions between NSCLC and Alzheimer’s disease patient-subgroup with at least 4 patients. Blue and red squares represent positive and negative relative risks respectively. **C)** Heatmap of the interactions between NSCLC and Alzheimer’s disease patient-subgroup with at least 4 patients with at least one drug associated in the same direction to all the patients within the same subgroup. Blue and red squares represent respectively positive and negative relative risks with shared drugs in the correct direction (at least a drug is associated in the same direction to all the patients within both subgroups in the case of positive interactions, and in opposite directions in the case of negative interactions).

To evaluate to what extent our measure of RR is able to reflect epidemiologically-defined comorbidities, we compared our DMSN with a undirected epidemiological network, the Phenotypic Disease Network (PDN), generated by Hidalgo et al. using the disease history of more than 30 million patients ^10^ (Methods). **Remarkably, our DMSN significantly recovers 20% of the PDN interactions (704 interactions, pval=0.00005, Fig. 3), as estimated by randomization (Methods)**. Then, in order to measure the similarity with a directed network, we compared our DMSN to the disease-pairs underlying temporal disease trajectories ^19^. We obtained a significant overlap between our pRR interactions and their disease-pairs (19, pval=0.014, Fig. S1). Interestingly, we additionally consider nRR interactions (as potential evidences of inverse comorbidity), which are not present in the epidemiological networks and constitute a new layer of knowledge.

**Figure 3.**
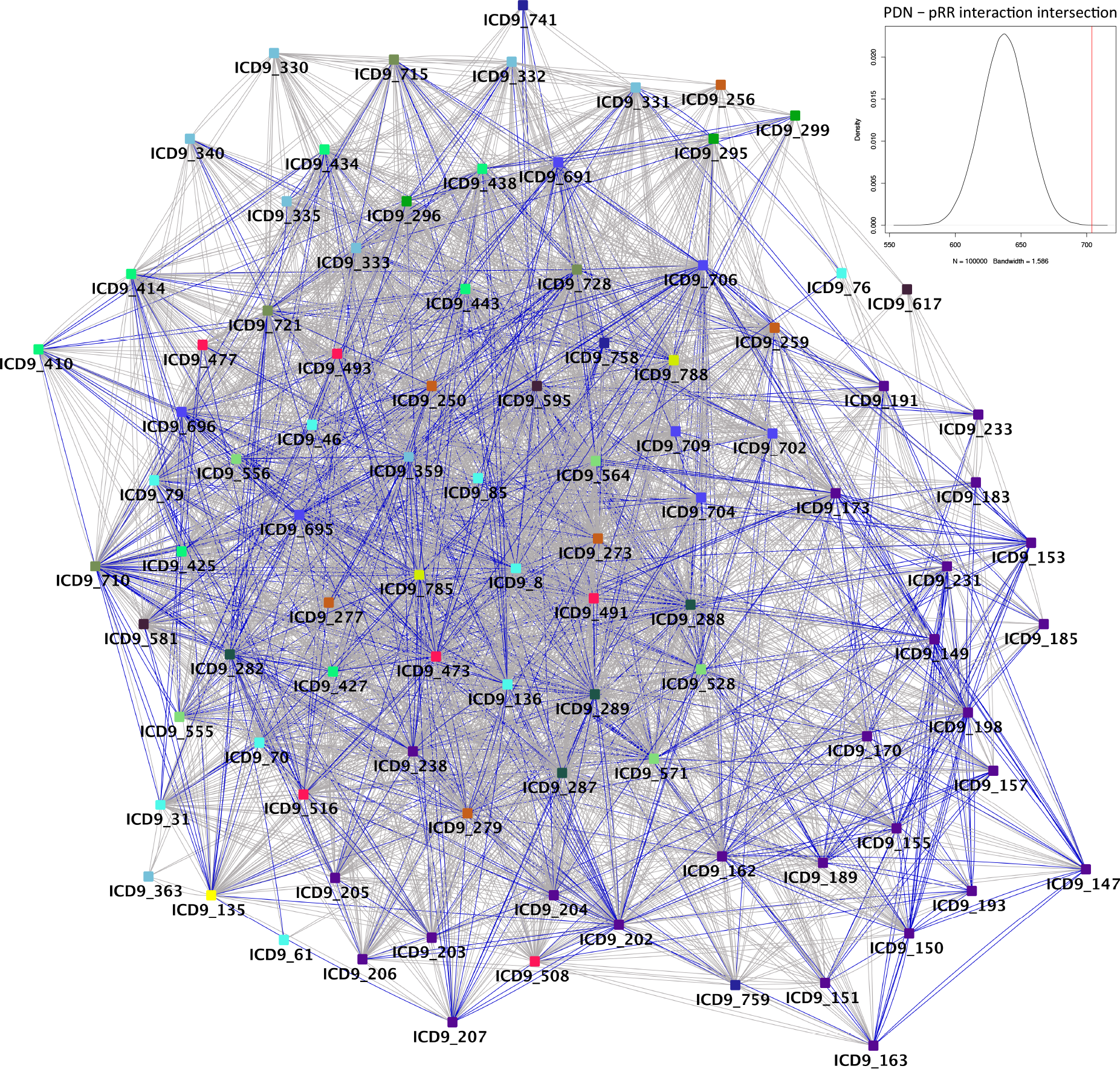
Phenotypic Disease Network - Diseases’ Molecular Similarity Network overlap.

The tendencies observed in the epidemiological studies and also in the DMSN indicate that at least a fraction of patients with one disease will have a higher probability of acquiring the second disease. Analyzing molecular similarities between patients suffering from the same disease, we observed different levels of heterogeneity (Fig. 4), which we define as the percentage ratio between observed vs. total number of possible intra-disease interactions. According to this definition, diseases with few intra-disease interactions, in which patients are less similar to each other, have higher molecular heterogeneity. The ICD9 categories “diseases of the skin and subcutaneous tissue”, “symptoms, signs, and ill-defined conditions” and “neoplasms” are the ones with the lowest molecular heterogeneity (Fig. 4). On the contrary, “mental disorders” and “diseases of the nervous system and sense organs” are the most molecularly heterogeneous ones (Table S1), potentially as a consequence of diagnostic methodologies. Such results denote that **high molecular and phenotypic heterogeneity might drive different comorbidity patterns in patients with the same disease**. We therefore generated subgroups of patients within each disease considering differentially expressed genes (Methods). In total we obtained 1,051 subgroups, 17% of them including patients from different studies (which constitutes 62% of the diseases with multiple studies), with an average number of 7.8 subgroups per disease. Interestingly, even large subgroups composed by 70 patients, share genes that are deregulated in the same direction in all the patients within it, supporting the reliability of molecularly defined patient-subgroups.

**Figure 4.**
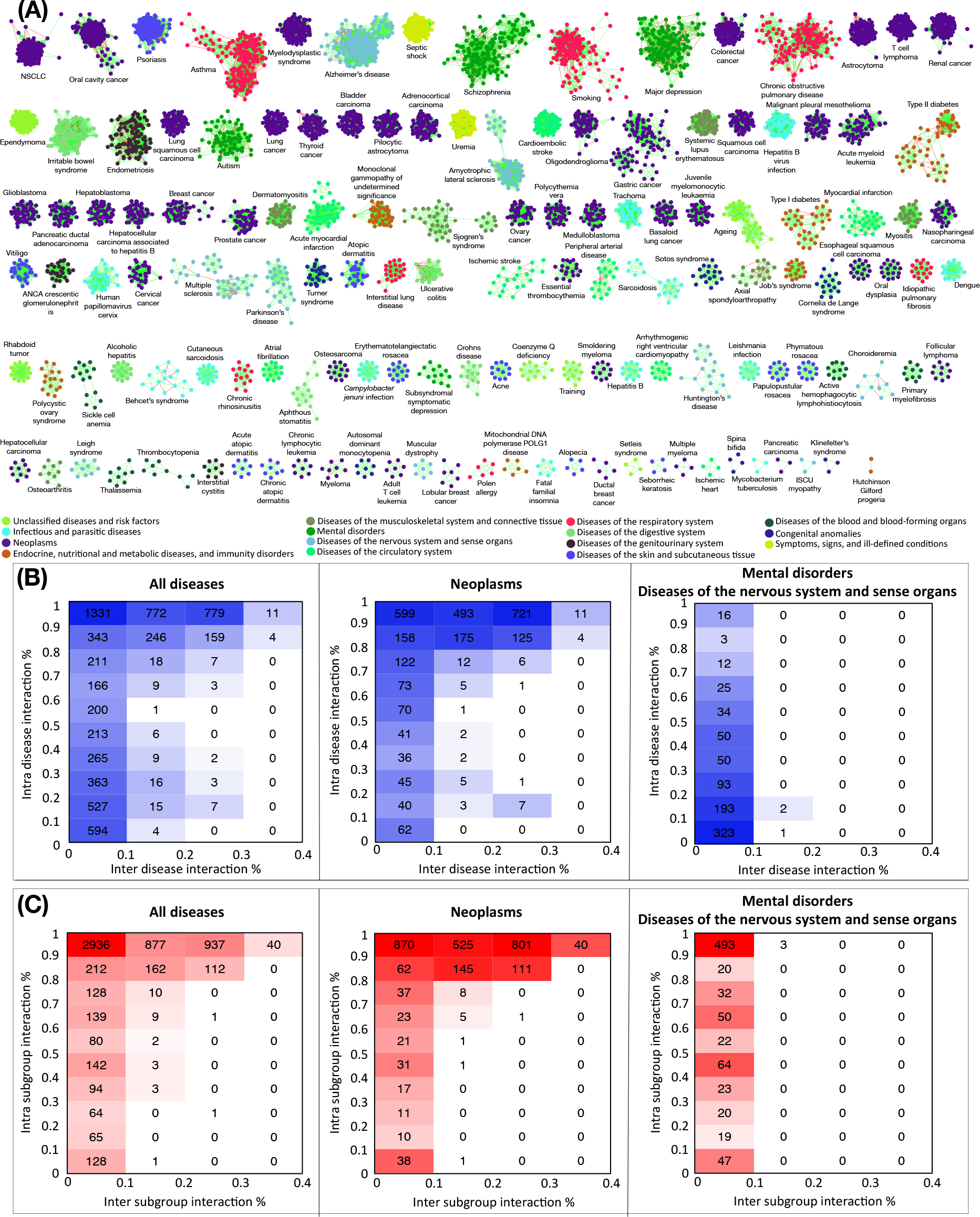
Intra-disease patient-patient interaction network and associated disease heterogeneity. **A)** Intra-disease patient-patient interaction network. Each node represents a patient. Green and red edges represent positive and negative interactions respectively. Nodes are colored based on the disease-group they belong to. Organic layout was used to represent the network ^51^. **B)** Patients’ intra-vs. inter-disease interaction percentages. Number of patients within each interval of intra-and inter-disease (or subgroup) percentage are indicated.

To quantitatively evaluate the heterogeneity of diseases and consistency of the subgroups, we calculated the intra-and inter-disease/subgroup interaction percentages for each patient (Fig. 4). Most patients presented a higher intra-subgroup than intra-disease interaction percentage. Such difference was especially higher in patients affected by more heterogeneous diseases, mental disorders and diseases of the nervous system, where inter-disease/subgroup interactions did not vary. In summary, disease-related molecular heterogeneity can indicate the presence of patient-subgroups, as observed in diseases such as diabetes ^20^ and different cancers ^21,22^. Regarding patient-subgroups, it is clear that, based on medical records, not all patients with a disease have a tendency to the same comorbidity patterns ^19^. **Defining these subgroups we provide the conceptual basis to design a clinical stratification based on patient-specific comorbidities.**

### Stratified Comorbidity Network

Considering interactions between patient-subgroups, we obtain a network with 1,051 nodes and 139,622 edges, which we call the Stratified Comorbidity Network (SCN). Exploring disease interactions at the more detailed level of patient-subgroups, we can potentially confirm relations observed between diseases, discover new relations not detected at the disease level, and also find comorbidities opposite to the ones described at the disease level.

We corroborated 95% (5,554/5,826) of pRR and nRR interactions detected in the DMSN. When considering patient-subgroups instead of diseases, we detected more than 8,000 new interactions in total. **Interestingly, 761 interactions revert the general trends observed at disease level, showing that not all the patients with a specific disease present the same comorbidity relations, as expected given the observed disease heterogeneity.**

To deepen the analysis of our SCN, we focussed on patient-subgroups composed of at least 4 patients with shared deregulated genes (Fig. S2). The resulting network comprises 182 subgroups and 1,624 interactions, corresponding to 385 interactions among 70 diseases. 39% of these disease-disease interactions were not present in our DMSN. Based on a curated list of PubMed papers, we observed that 55% of such patient-subgroup interactions have been previously described by epidemiological studies (Fig. 5, Supplementary Text). For example, we observed a higher than expected risk of developing AD in a subset of smokers, a tendency which was previously suggested in the literature ^23^.

**Figure 5.**
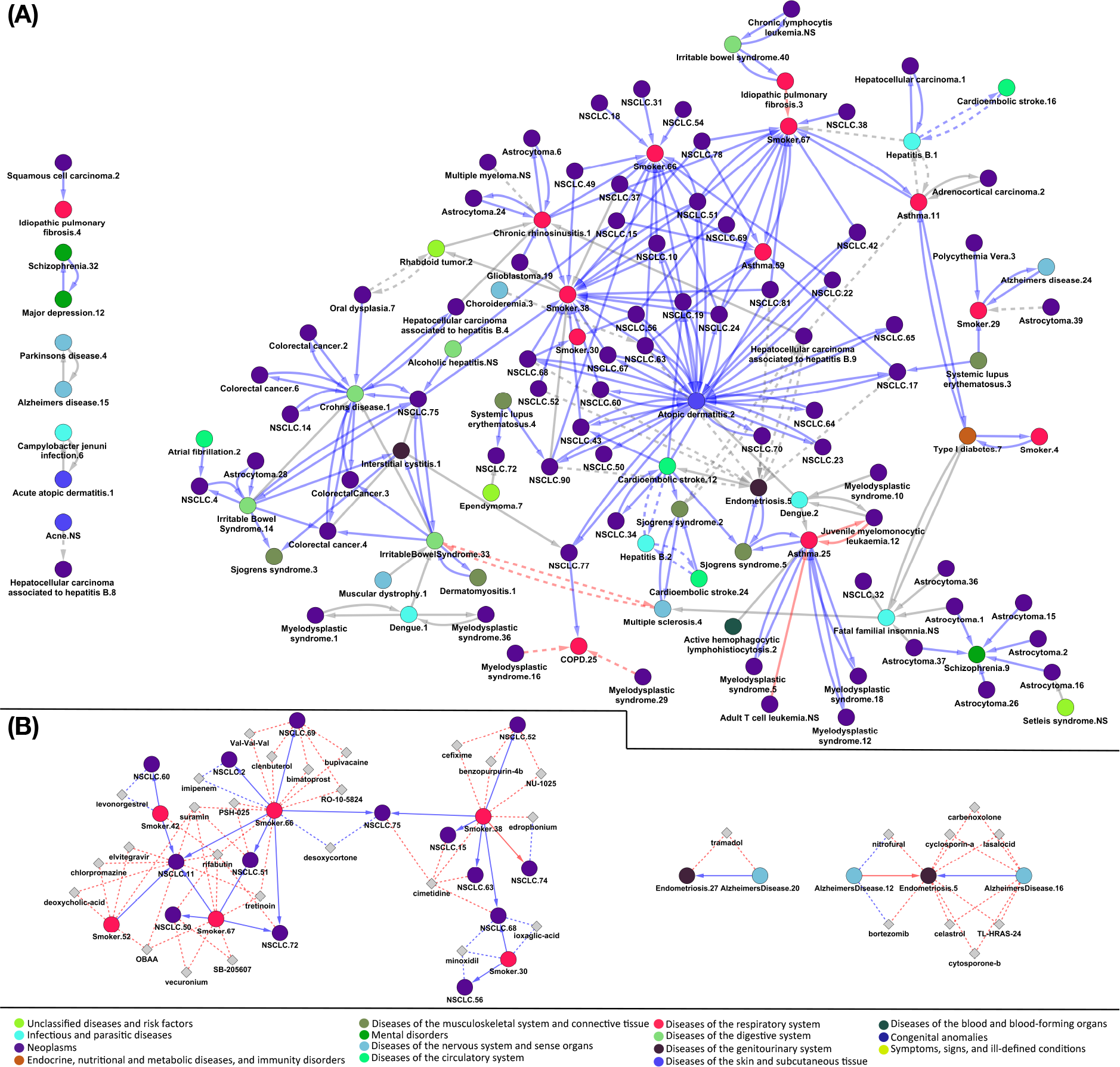
Size 4 patient-subgroups with shared genes (a) and drugs (b & c). **A)** Each node represents a patient-subgroup, colored based on the disease-group they belong to. Solid and dashed lines represent positive and negative relative risk interactions. Blue, red and grey colored interactions represent interactions matching, opposing and not previously described in epidemiological data respectively. **B & C**) Circle and diamond nodes represent patient-subgroups and drugs respectively. Blue and red colored edges represent positive and negative interactions respectively. Solid lines denote relative risk interactions while dashed lines denote subgroup-drug interactions.

These results show that our networks (DMSN and SCN) derived from expression data recapitulate many previous epidemiological results, showing their applicability in the discovery of novel direct and inverse comorbidity relations between diseases. Importantly, we go beyond disease definitions and find positive and negative comorbidity relationships between specific patient-subgroups.

### Association of differential expression profiles to known drug effects

Since gene expression can be altered by drug intake, we investigated if any of the observed interactions could be related with the effects of drugs on gene expression patterns as previously done by Jahchan et al. ^24^. To this end, we compared patients’ differential expression profiles with those reported in the LINCS L1000 library (Methods). We added LINCS drugs as nodes in our SCN and investigated whether specific patient-subgroups have any common drug associations. If the changes generated by the drug were similar to the ones observed in specific patient-subgroups, we could surmise that the drug is responsible for such patterns. On the other hand, if the changes were opposite to the ones observed in patient-subgroups, then this would suggest that the drug could serve to treat those patients specifically, opening the way to drug repurposing ^24,25^. Our results show that patients within each subgroup had significantly more common drugs associated to them than expected by chance (Methods, Fig. S3.).

For in-depth analysis of the possible relations between diseases and drugs, we restricted our network to subgroups composed of at least 4 patients with expression profiles correlating with a given drug. From the resulting network, we obtained 152 cases where subgroups from 2 different diseases presented both negative and positive RR relations while being associated with the same drug. For example, we obtained 20 pRR interactions between 6 smoker and 14 NSCLC subgroups, and one nRR interaction, with edrophonium being identified as a potential player in the nRR interaction (Fig. 5). These results suggest that, despite the well known increased risk of developing lung cancer in smoking patients, a small subset of smokers might be at a lower risk of developing the disease due to their specific molecular characteristics.

Another noteworthy example is the case of AD and endometriosis, which are directly comorbid based on our disease level network. In the SCN, we obtain two pRR and one nRR between 3 AD and 2 endometriosis subgroups (Fig. 5) and observe interactions with bortezomib, which suggests that this drug can potentially increase the risk of AD, as well as protect against endometriosis. Interestingly, bortezomib is currently being explored as a therapeutic option for endometriosis ^26^.

Strikingly, **we identified different molecular mechanisms potentially involved in the comorbidity between specific patient-subgroups in the same pairs of diseases** (Fig. 2).

For example, focusing on the AD - NSCLC inverse comorbidity relation, we detected 142 nRR interactions between patient-subgroups. Selecting only those interactions with an associated drug, this number decreased to 18. Interestingly, several drugs targeting different molecular mechanisms were detected in those nRR interactions, with MST312 (telomerase inhibitor), tacrolimus (immunosuppressive agent), and Antimycin A (antibiotic) among others, suggesting that different molecular mechanisms might explain the same relationship between diseases.

<u>In summary, the use of the SCN filtered by shared drugs allows a deeper analysis of the molecular processes potentially involved in comorbidity. This approach is especially interesting in those cases in which a set of patients present comorbidity relations opposite to the ones observed at the disease-level.</u>

### Patients’ comorbidity profiles

The final application of the presented approach is to develop a methodology that could be used to identify the most probable comorbidities each specific patient is likely to develop. To this end, for each patient, we ranked the diseases from the most probable to the least based on patients’ molecular similarities (Methods), associating LINCS drugs to the comorbidity risks. Finally, we looked for examples where a patients’ first-line-treatment might be causative of increasing the risk of developing the most probable secondary diseases (Fig. S4). As an example, we detected one AD patient connected to rivastigmine (an inhibitor of acetylcholinesterase used for AD treatment) at a significant RR of developing muscular dystrophy. Since 6 out of 7 patients with muscular dystrophy are positively connected to rivastigmine, it could be speculated that treating this specific AD patient with rivastigmine would increase his/her risk of developing muscular dystrophy, suggesting that alternative treatments should be sought.

This approach can be extended systematically to all the other patients and diseases, but each case requires a careful analysis of the relation with drugs. In addition to the proof of principle results reported in this paper, we make all the generated results accessible to the research community through the Disease PERCEPTION portal (http://disease-perception.bsc.es/), which allows exploration of the Diseases’ Molecular Comorbidity Network and the Stratified Comorbidity Network (Fig. 6).

**Figure 6.**
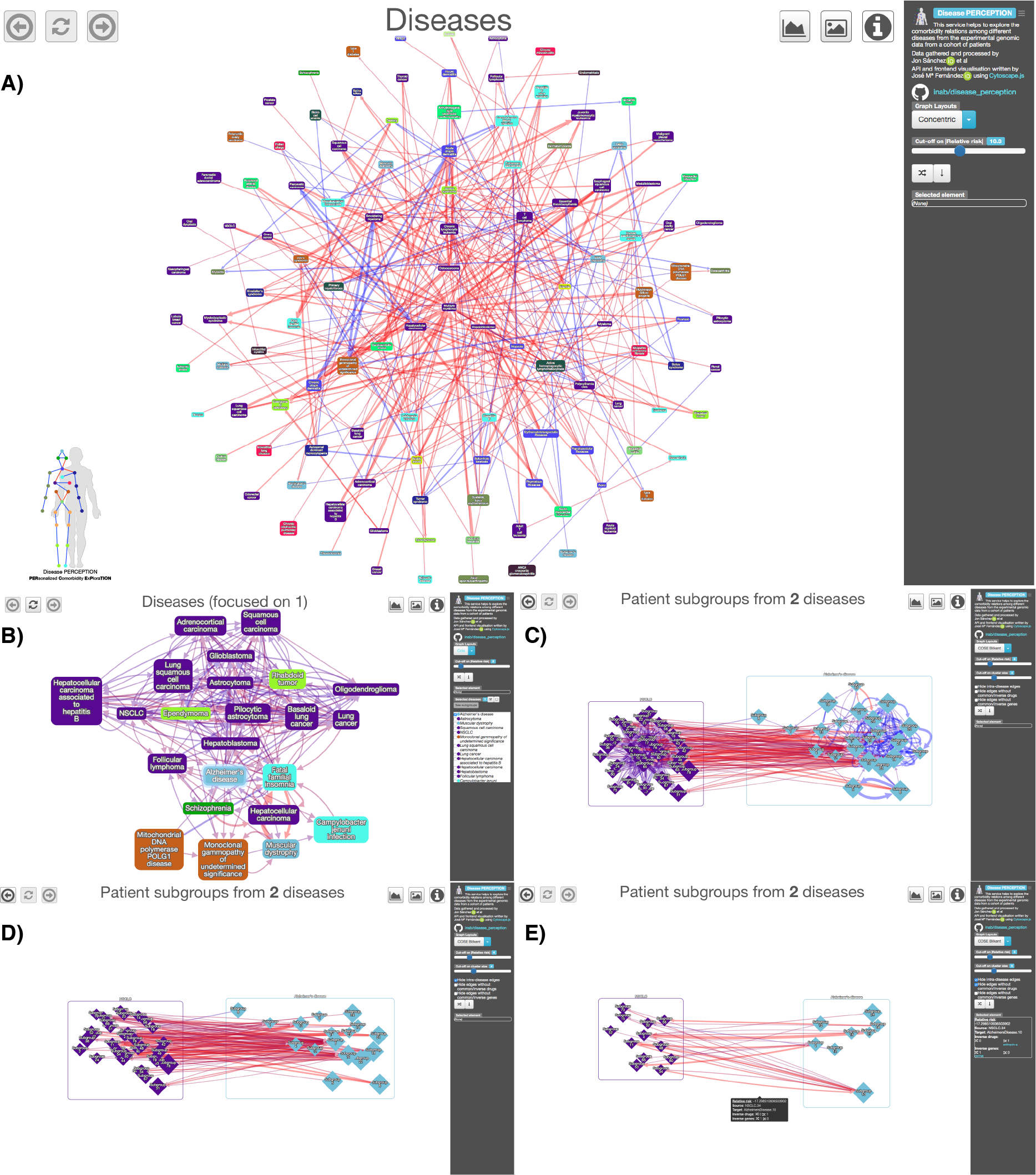
The Disease PERCEPTION portal. Through this user-friendly and programmatically accessible portal, the user can visualize comorbidity relations at the disease and patient-subgroup levels. Moreover, users can extract patient-subgroup information, filtering by subgroup size, intra-subgroup connectivity, as well as by shared drugs and/or genes. Genes and drugs in the networks are hyperlinked to databases, facilitating an interactive exploration of the molecular basis of each connection. **A)** Disease network view. Each node represents a disease, coloured based on the disease category they belong to. Blue and red edges denote positive and negative relative risk (pRR, nRR) interactions. Relative risk cut-off can be modified. **B)** Alzheimer’s disease neighbours view. Desired diseases can be selected to show their patient-subgroups. **C)** Alzheimer’s disease and non-small cell lung cancer patient-subgroups with >4 patients per subgroup. **D)** Same as C) excluding intra-disease interactions. **E)** Same as C) showing only patient-subgroup interactions with shared drugs. Selecting edges of interest displays genes and drugs potentially involved in the selected interactions.

## Discussion

In this study we produced networks that are predictive of comorbidity risk at disease and patient level by estimating patient-patient similarities based on differential gene expression information between cases and controls We identified transcriptional heterogeneity in different diseases, which could reflect the presence of molecular disease subtypes ^20^ or low-specificity in the diagnostic procedures (indeed, conditions diagnosed using more accurate methods like biopsies - neoplasms - show lower transcriptomic heterogeneity compared to others, e.g. Central Nervous System disorders, based on neurocognitive evaluation). Differences between patients can be due to genetics and the environment ^27^, more explicitly living conditions, food and drug intake ^28^. Despite all these factors contributing to the heterogeneity of the expression profiles, the similarities measured as pRR between diseases are strong enough to reveal significant overlaps with relations previously observed at epidemiological level. Furthermore, the presented molecular analysis allows the detection of nRR, that are not reported in the large epidemiological studies, but are well known at the level of specific population studies, including the ones between Central Nervous System and cancer ^29,30^.

As a demonstration of the potential of this approach, the selection of the positive and negative RR interactions in the SCN based on shared deregulated genes allows for a deeper analysis of the molecular bases of comorbidities. For example, a potential driver of the increased risk of developing AD in smoking patients detected in our study (F13A1) encodes the coagulation factor XIII A subunit, which has been described to be up-regulated in smokers ^31^ and also in AD patients, where it has been proposed as a serum biomarker that correlates with PiB-PET data as an early diagnostic method of the disease ^32^.

Further, we found patient-subgroup comorbidities that defy the ones described at the disease level. For example, we identify a subgroup of smokers who might be protected against the development of NSCLC. Interestingly, using LINCS for relating drugs to expression profiles, a cholinesterase inhibitor (edrophonium) is found to produce a transcriptional signature compatible with the expression changes observed in the smokers subgroup, which are also opposite to the ones observed in the NSCLC patient-subgroup. Remarkably, acetylcholinesterase inhibitors have been shown to reduce nicotine dependency ^33^ while contributing to lung cancer progression ^34^, suggesting that the intake of these drugs to reduce nicotine dependency by this subgroup of smokers might increase their risk of developing NSCLC. An additional case of inverse comorbidity is the one between AD and endometriosis, in which the drug that increases the risk of AD may at the same time reduce the risk of endometriosis. Interestingly, the drug bortezomib might generate expression changes similar to the ones observed in AD by inhibiting the proteasome and thus mediating amyloid-beta neurotoxicity ^35^. This drug has been proposed as a potential novel therapeutic strategy for the treatment of endometriosis ^26^.

The importance of a personalized approach to comorbidity relations is evidenced by the association of different pairs of AD and NSCLC patient-groups to different drugs, which suggests different molecular mechanisms underlying the same protective effect. Interestingly, the altered molecular mechanisms detected by our analysis have been previously described in both diseases separately. Telomerase inhibition shows a strong antiproliferative effect on lung cancer ^36^ and, at the same time, a significantly accelerated rate of telomere shortening has been described in AD patients ^37^. The use of tacrolimus was shown to attenuate cognitive deficits and oxidative stress in rats with induced AD type dementia ^38^ while increasing the risk of developing solid tumours after liver transplant ^39^.

<u>Taken as a whole, our results suggest that investigating expression profiles could be a useful tool that allows the detection of different processes potentially driving comorbidities between pairs of diseases. The comparison of expression profiles as indicators of physiological states serves as an initial quantification of how different patients with the same disease have different comorbidities driven by different molecular processes. Our approach suggests that it might be possible to reverse-engineer the network to deduce genes or pathways likely to underlie the observed relationships, identifying comorbidity biomarkers.</u>

In conditions that are diagnosed based on biopsies, transcriptomic data will be available very soon at a limited cost. The characterization of patients’ molecular phenotypes, through transcriptomics, proteomics or other experimental techniques, will allow a deeper study of complex comorbidity patterns and mechanisms, beyond the statistical picture traditionally provided by epidemiological approaches. Indeed, the consistency of the results for different tissues and diseases suggests that the molecular basis of comorbidities have a systematic character, and profiling patients’ blood samples in the future could be used to produce comorbidity risk profiles, as suggested in other scenarios ^40,41^. Understanding and managing multi-morbidity has been identified as a priority for global health research ^42^. We argue that a person’s molecular profile can be used as a predictor of disease comorbidity risk and as a key component in a personalized comorbidity management strategy.

## Methods

### Gene expression analysis

Gene expression raw data (CEL files) were downloaded from the Gene Expression Omnibus (GEO, GSE* files http://www.ncbi.nlm.nih.gov/geo) and ArrayExpress (EMTAB* files https://www.ebi.ac.uk/arrayexpress/) for 133 diseases and 3 risk factors, including 186 datasets (Table S2). Studies undertaken on HG U133Plus2 Affymetrix microarray platform were selected to allow using the frozen Robust Multiarray Analysis (fRMA) normalization method ^43^ and reduce the bias due to inter-platform differences. The linear regression model provided by the LIMMA package was used to identify differential gene expression ^44^, comparing each sample case with all the control samples from the same study (from now on denoted as patient).

Interactions between patients within the same disease (using the same threshold used in previous studies ^16,18^) were calculated, varying the number of genes selected as significantly differentially expressed, demonstrating that the number of detected significant interactions between patients increases while increasing the number of selected genes (Fig. S5). The Top 500 up-and down-regulated genes were selected as significantly differentially expressed for posterior analyses based on the t-values provided by LIMMAs differential gene expression analysis.

### Patient-patient interaction analysis

Following the strategy reported by Ibañez et al., 2014 ^16^, overlaps between pairs of patients were assessed by one-tailed Fisher’s exact tests on lists of DEGs. Two patients are positively connected if they present significant overlaps between genes deregulated in the same direction (both up-and down-regulated). On the other hand, two patients are negatively connected if they present significant overlaps between genes up-regulated in one patient and down-regulated in the other one, and vice versa. If both types of significant overlaps are detected in the comparison (e.g. significant overlaps between genes up-and down-regulated in both patients and between genes up-regulated in one patient and down-regulated in the other one), there is no link between those patients.

1,000 permutations were conducted randomly selecting 500 genes as up-and down-regulated. The Fisher’s exact test threshold for the detection of significant overlaps was varied to establish a value for which no significant interactions were detected in the random cases (FDR≤0.0005, Fig. S6).

### Patient clustering

Same disease patients were clustered based on their discretized differential gene expression information into disease subgroups denoted from now on as patient-subgroups. The optimal number of clusters within each disease was obtained using the Silhouette method ^45^, where k-means analyses were conducted using Hartigan and Wong’s algorithm ^46^ varying the number of clusters from 2 to the total number of patients within the disease. The number of clusters with the highest silhouette score was selected as the optimal number of clusters.

For each disease we extracted the total number of genes commonly deregulated in the same direction in all the patients within each subgroup. Then, we shuffled 1,000 times the patients among the subgroups from the same disease and calculated the total number deregulated genes. Only those patient-subgroups from diseases with a total number of shared genes higher than the expected by chance were selected as true patient-subgroups.

Additionally, we extracted the number of patient-subgroups with shared genes, selecting those subgroups with at least one gene up-and one gene down-regulated in all the patients composing the subgroup. This number was compared to the values expected by chance, shuffling 10,000 times for each disease the patients and extracting the number of subgroups with shared genes. Most patients within subgroups shared at least one gene deregulated in the same direction, more than expected by chance (98% vs. 79%) Fig. S7. The same was done with the drugs, estimating the number of subgroups with at least one drug connected with the same sign to all the patients within a subgroup.

To deepen our analysis of the molecular bases of comorbidity relations between patient-subgroups, we filtered those subgroups with at least 4 patients and shared molecular alterations, in terms of commonly up-and down-regulated genes (at least one gene up-and one gene down-regulated in all the patients), selecting only those interactions with overlapping genes deregulated in the same direction in both patient-subgroups for positive interactions, and in opposite directions for negative interactions.

### Disease heterogeneity analysis

Looking at the patient interaction network defined previously, we estimated whether patient clusters were consistent with grouping by disease and by patient-subgroups. The agreement was assessed based on intra-and inter-disease interaction percentages for each disease separately.

### Relative risk estimates

For each pair of diseases we consider a contingency table counting the number of positive interactions connecting patients from the two diseases and the ones connecting one of the diseases with other ones.

|  | **Disease B** | **No disease B** | **Total** |
| --- | --- | --- | --- |
| **Disease A** | $N_{ab}$<br>(interactions of interest) | $N_{anb}$<br>(other interactions) | $T_a$ |
| **No disease A** | $N_{nab}$<br>(interactions of interest) | $N_{nanb}$<br>(other interactions) | $T_{na}$ |

We can then define the proportion of interactions connecting patients from the two diseases of interest to the total number of interactions of disease A (Pexposed) and the proportion of total number of interactions connecting patients of disease B and diseases other than A, compared to the total number of interactions outside of disease A (Pnonexposed). Positive interactions between patients are considered as interactions of interest, merging both negative and no-interactions as the other interactions.

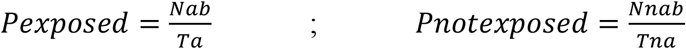

These quantities allow us to define relative risk (RR) for each pair of diseases, according to the following formula:

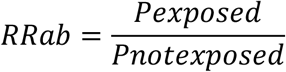

Repeating the same procedure using negative interactions (considering positive interactions as no interactions) we similarly define negative Relative Risks (nRR).

95% confidence intervals were calculated for diseases, patient-subgroups and patient-disease relations using the following formula.

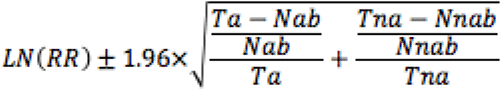

### Comparison with epidemiological networks

To validate our results regarding comorbidities, we compared them with the ones obtained at an epidemiological level by Hidalgo ^10^ and Jensen ^19^. Hidalgo et al. used ICD9 disease codes to associate patients to diseases generating a disease-disease network called PDN. We therefore started by grouping our patients using the same disease taxonomy and calculated relative risks between ICD9 codes as previously done at the disease level, reducing the confidence interval to 99% as in their analysis. To verify the significance of the overlap between our relative risks and protections with the ones detected by Hidalgo et al, we conducted 100.000 randomizations generating random interactions between the common set of 94 ICD9s. The same was done to compare our network with the disease trajectories, associating in this case our patients to ICD10 codes.

### L1000 LINCS analysis

The t-values obtained for each patient when compared against all the control samples in each disease were used as gene expression signatures of the patients, and compared against the LINCS L1000 library (http://www.lincscloud.org/), as performed previously. The LINCS L1000 library is a large catalogue of gene expression signatures in cancer cell lines induced by drug treatment or gene knockdown ^47^.

From the L1000 library drug induced expression signatures were obtained from experiments in which the transcriptional state of the cell is measured before and after the treatment with the drug. This allows to study the transcriptional effect of the drug. In order to obtain consensus expression signatures for each drug, a differential expression analysis was performed on control vs. treated cells using limma ^44^.

In the LINCS L1000 data, all the wells in which the same drug was used were considered as treated samples. All the DMSO treated wells from all the plates with at least one treated well were considered as untreated controls. The plate in which the drug was tested was taken as a covariate in the expression analysis. As only a type of cell line is used in each plate, using this covariate we take into account the technical batch due to different plates and the biological variability due to different cell lines. Different drug concentrations and exposure time to the drug were not taken into account. In the LINCS L1000 data some drugs (pert_iname) are represented by different molecules, (pert_id) usually from different vendors. In these cases, we obtained the pert_id associated signatures, that is, associated to the molecule, and a consensus signature in which all the pert_id corresponding to the same drug were considered. In this last case, the pert_id was also taken as a confounding variable.

The t-moderated statistic was used as a measure of the expression of the gene. It was preferred over the logFC because the t statistic takes into account the sampling variance. However, both statistics were highly correlated in all the signatures tested.

In order to measure the similarity of each patient signature to a given drug signature, the enrichment of the top 250 up-regulated and down-regulated genes by the drug was determined in the patient signature using a pre-ranked GSEA.

The fgsea R package was used ^48^. A consensus Enrichment Score (ES) was obtained subtracting the ES values of the DN signature from those ES of the UP signature.

### New patient classification and comorbidity prediction

Each patient of the study was classified into their corresponding disease and patient-subgroup using a leave one out approach, comparing the patients’ differential gene expression profile with the ones of each other patient (up-regulated genes were denoted with 1s, down-regulated ones with -1s and all the other ones with 0s) using euclidean distances. TP, TN, FP and FN values were calculated for each disease, and for each patient-subgroup within the same disease. Precision, recall and specificity values were calculated (selecting the same number of TP-FP and TN-FN) and compared to random expectation, shuffling 10,000 times the gene expression values across patients.

Then, two new AD and NSCLC datasets, analyzed using the same microarray platform (HG U133 Plus 2), were downloaded from GEO (GSE84422 and GSE27262), with 17 and 25 patients respectively, and classified into one of our 136 diseases and risk factors based on their differential gene expression profiles.

### Personalized comorbidity profiles

For each patient, based on patients’ molecular similarities, we calculated the pRR and nRR of developing each of the analyzed diseases as done before with diseases and patient-subgroups, producing a ranked disease list from the most probable to the least. Then, for each disease we added LINCS drugs and ranked them from the one similar to most patients to the one similar to the least, highlighting the first-line-treatments (https://www.vademecum.es). As a final step, we look for examples where a patients’ first-line-treatment might be causative of increasing the risk of developing the most probable secondary diseases (Fig. S4), this is, drugs that are positively connected to most patients of the secondary diseases.

### Disease PERCEPTION portal

The portal is composed by a database loader, a SQL database, a REST API and a web frontend. The tabular data and the source code of the database loader, REST API and web frontend are available at the GitHub project https://github.com/inab/disease_perception.

The database loader is written in Python 3.5, and it uses pandas ^49^ and SQLite to prepare a SQLite database instance. The SQL database is composed by 16 tables, with the disease groups, diseases, patient subgroups, patients, studies, genes, drugs and their relationships.

The data loaded comes from all the results consolidated from the analyses previously described.

The REST API is written in Python 3.5, and it uses Flask, Flask-RESTPlus and Flup. It is available at http://disease-perception.bsc.es/api/, and it is documented using OpenAPI.

The Disease PERCEPTION web frontend is written in Javascript ES7/ES2016, and it uses Cytoscape.js ^50^, the external layout plugins COLA, COSE-Bilkent, Dagre and Klay, JQuery, Bootstrap, Tippy and Popper. It is built using yarn, babel and webpack, as it is described in its documentation on the GitHub repository.

## Acknowledgments

We thank Anaïs Baudot (Université d’Aix-Marseille), Eduard Porta (Barcelona Supercomputing Center), Alba Jene (Barcelona Supercomputing Center) and Davide Cirillo (Barcelona Supercomputing Center) for critical reading of the manuscript and helpful discussions.

## Funding

This work was supported by a PhD Fellowship (BES-2016-077403) and funded by the Spanish Ministry of Economics and Competitiveness (BFU2015-71241-R). The Coordination Node led by Dr. Alfonso Valencia at the Barcelona Supercomputing Center (BSC) is a member of the Spanish National Bioinformatics Institute (INB), ISCIII-Bioinformatics platform and is supported by grant PT17/0009/0001, of the Acción Estratégica en Salud 2013-2016 of the Programa Estatal de Investigación Orientada a los Retos de la Sociedad, funded by the Instituto de Salud Carlos III (ISCIII) and European Regional Development Fund (ERDF). Rafael Tabarés-Seisdedos was supported in part by grantnumber PROMETEOII/2015/021 from Generalitat Valenciana and the national grant PI17/00719 from ISCIII-FEDER.

## Author contributions

A.V., V.P., D.J., and J.S. designed all experiments. J.S. and H.T. performed the experiments. F.A. and R.T. provided technical advice. J.M.F. and S.C. generated the Disease PERCEPTION portal. J.S., V.P., and A.V. wrote the manuscript. All authors discussed the results and commented on the manuscript.

## Competing interests

The authors declare no competing interests.

## Data and material availability

All data needed to understand and assess the conclusions of this research are available in the main text, supplementary materials and Disease PERCEPTION portal.

